# Sulforaphane Alters the Acidification of the Vacuole to Trigger Cell Death

**DOI:** 10.1101/371534

**Authors:** Alexander Wilcox, Michael Murphy, Douglass Tucker, David Laprade, Breton Roussel, Victoria Hallisey, Christopher Chin, Abraham Brass, Nicanor Austriaco

## Abstract

Sulforaphane (SFN) is a compound [1-isothiocyanato-4-(methylsulfinyl)- butane] found in broccoli and other cruciferous vegetables that is currently of interest because of its potential as a chemopreventive and a chemotherapeutic drug. Recent studies in a diverse range of cellular and animal models have shown that SFN is involved in multiple intracellular signaling pathways that regulate cell death, cell cycle progression, and cell invasion. In order to better understand the mechanisms of action behind SFN-induced cell death, we undertook an unbiased genome wide screen with the yeast knockout (YKO) library to identify SFN sensitive (SFN^S^) mutants. Our mutants were enriched with knockouts in genes linked to vacuolar function suggesting a link between this organelle and SFN’s mechanism of action in yeast. Our subsequent work revealed that SFN increases the vacuolar pH of yeast cells and that varying the vacuolar pH can alter the sensitivity of yeast cells to the drug. In fact, several mutations that lower the vacuolar pH in yeast actually made the cells resistant to SFN (SFN^R^). Finally, we show that human lung cancer cells with more acidic compartments are also SFN^R^ suggesting that SFN’s mechanism of action identified in yeast may carry over to higher eukaryotic cells.

## Introduction

The consumption of broccoli and other cruciferous vegetables belonging to the *Brassica* family has been shown to have protective effects against several types of cancer, including prostate, breast, colon, and lung cancer. [1, 2] Though these plants contain a diverse range of metabolites and antioxidants, the chemical agents believed to be responsible for these effects are the naturally occurring organosulfur compounds called isothiocyanates (ITCs; R-N=C=S). [3, 4]. These molecules are the products of the reaction of plant glucosinolates with myrosinase, an enzyme released by the disruption of plant tissues.

Studies undertaken during the past two decades have reported that the isothiocyanate in cruciferous vegetables primarily responsible for their chemopreventive effects is the isothiocyanate called sulforaphane (SFN; 1- isothiocyanato-4-(methylsulfinyl)butane). [5] Numerous experiments have shown that SFN can defend healthy cells against chemical and radiation induced carcinogenesis and can inhibit the proliferation, migration, and survival of tumor cells. [6, 7] There is also extensive evidence that reveals that SFN is a chemoprevention agent against cardiovascular diseases, neurodegenerative diseases, autism, and diabetes. [8-10]

Sulforaphane affects many molecular targets in cellular and animal models. However, its cytoprotective function has been attributed primarily to its diverse abilities. These include SFN’s abilities to inhibit phase 1 metabolizing enzymes (mostly cytochrome P450); to alter the localization of the transcription factor Nrf2 so that it can enter the nucleus to regulate the basal and inducible expression of a multitude of antioxidant proteins, detoxification enzymes, and xenobiotic transporters; and to suppress pro-inflammatory responses within the cell. [4] SFN is also known to inhibit histone deacetylase, which could explain its ability to induce cell cycle arrest and apoptosis, and to regulate different microRNAs. [11-13] Finally, there is data that suggests that SFN can trigger cell death by upregulating caspases and downregulating anti-apoptotic factors. [14-16]

In order to better understand the mechanisms of action of SFN in eukaryotes and to possibly uncover novel ones, we undertook an unbiased genome wide screen with the *Saccharomyces cerevisiae* knockout (YKO) library, a collection of 4,775 individual yeast strains, each of which contains a deletion of a single non-essential yeast ORF, to identify mutations that affect the cell’s sensitivity to SFN. The YKO collection has been used extensively over the past decade to identify the mechanisms of actions of a wide range of small molecules and drugs. [17, 18] Our screen uncovered numerous SFN^S^ mutants. Notably, they were enriched with knockouts in genes linked to vacuolar function suggesting a link between this organelle and SFN’s mechanism of action in yeast. Our subsequent work revealed that SFN increases the vacuolar pH of yeast cells and that varying the vacuolar pH can alter the sensitivity of yeast cells to SFN. In fact, several mutations that lower the vacuolar pH in yeast actually made the cells resistant to SFN. Finally, we show that human lung cancer cells with decreased endosomal pH are also SFN^R^ suggesting that SFN’s mechanism of action identified in yeast may carry over to higher eukaryotic cells.

## MATERIALS AND METHODS

### Yeast Strains and Growth Conditions

All experiments were done with isogenic *Saccharomyces cerevisiae* strains in either the BY4742 (MATα *his3∆1, leu2∆0, lys2∆0, ura3∆0*) or the PSY316AR (MATα RDN1::ADE2 his3- 200 leu2-3,112 lys2 ura3-52) backgrounds. For all the experiments described in this paper, cells were cultured and treated using standard yeast protocols. [19] Unless noted otherwise, all drugs and reagents were purchased from SIGMA-Aldrich. Isothiocyanates were resuspended in acetonitrile as a solvent.

### Spot Assay

Seed cultures of the BY4742 and PSY316AR yeast strains were grown overnight in YPD. Each strain was diluted to an OD_600_ of 0.1 in fresh YPD and grown for at least 2 doublings (~5 hours). After the yeast strains entered log phase (OD_600_ ~0.4-0.8), sulforaphane (LKT Laboratories), BITC, or PEITC was added to the cultures at the indicated concentrations with the solvent, acetonitrile alone, as the no-drug control. Following the indicated incubation times, cells were removed, spun, washed, and diluted. For each strain, a series of 10-fold dilutions was then prepared in water over a range of concentrations from 10^−1^ to 10^−5^ relative to the initial culture. Spots of 5μl from each dilution series were then plated on the indicated media and cultured at 30^°^C for 2 days. All spot assays were repeated at least three times and a representative experiment is shown.

### Liquid Viability Assay

Seed cultures of each yeast strain were grown overnight in YPD. Each strain was diluted to an OD_600_ of 0.1 in fresh YPD and grown for at least 2 doublings (~5 hours). After the yeast strains entered log phase (OD_600_ ~0.4-0.8), SFN was added at the indicated concentrations. Cell viability was measured at the indicated time points following drug addition using a Nexcelom Vision Cell Analyzer with propidium iodide as a vital stain (1μg/ml). Statistical significance was determined with the Student’s t-test, using Graph Pad Prism 6. By default, one asterisk is p<0.05; two asterisks is p<0.01; three asterisks is p<0.001; and four asterisks is p<0.0001.

### Genetic Screen for SFN^S^ Mutants

Seed cultures of individual yeast strains from the BY4742 knockout library (Dharmacon Yeast Knock Out MATalpha Collection) were grown overnight at 30°C in 96-well plates in complete synthetic defined (SD) media. A 10μl aliquot of each culture was then transferred to a well of two different sets of 96-well plates, each of which contained 150μl fresh complete SD media. Cells were allowed to reach exponential phase (OD_600_ ~0.4-0.8). SFN was then added to one of the sets of 96-well plates to a final concentration of 200μg/ml. Relative growth for SFN^S^ mutants was determined by visual inspection of the wells, comparing wells with drug with wells without drug, after they had been cultured at 30^°^C for 2 days.

### Functional Gene Ontology Annotation

The Cytoscape 2.8.3 plugin BiNGO (v2.44) was used to identify enriched biological processes in the SFN^S^ mutant pool after Benjamini & Hochberg false discovery correction for multiple hypothesis testing.

### Confocal Imaging of Yeast Cells

BCECF-AM (Molecular Probes, Eugene, OR) staining was performed as described [20] with the following modifications: Seed cultures were grown overnight in YPD. Each culture was then diluted to an OD_600_ of 0.1 in fresh YPD and grown for at least 2 doublings (~5 hours). Once the cells were in log phase, sulforaphane, BITC, or PEITC were added to the cultures at the indicated concentrations with the solvent, acetonitrile alone, as the no-drug control. After they were allowed to grow at 30°C for an additional 18 hours, cells were harvested, washed, and resuspended in an equivalent amount of APG (a synthetic minimal medium containing 10mM arginine, 8mM phosphoric acid, 2% glucose, 2mM MgSO4, 1mM KCl, 0.2mM CaCl2, and trace minerals and vitamins titrated to pH 7.0 with KOH and 10mM MES). Two 200μl aliquots of each yeast culture were then transferred to a 96-well plate. They were incubated with 50 μM BCECF-AM at 30°C for 30 min, washed, and resuspended in APG medium to be imaged. Images were captured with a Zeiss LSM 700 Laser Confocal Microscope (Zeiss, Thornwood, NY), and processed using the Zen 2009 software package.

### Assay for the Measurement of Yeast Vacuolar pH

Seed cultures of each yeast strain were grown overnight in YPD. Each strain was diluted to an OD_600_ of 0.1 in fresh YPD and grown for at least 2 doublings (~5 hours). After the yeast strains entered log phase (OD_600_ ~0.4-0.8), cells were spun down and resuspended in APG media titrated to pH 3, 5, 7, 9, or 11. After an additional hour of growth in this media, the cells were incubated with 50μM BCECF-AM at 30°C for 30 min, washed, and resuspended in APG medium to be imaged. Images were captured with a Zeiss LSM 700 Laser Confocal Microscope (Zeiss, Thornwood, NY), and processed using the Zen 2009 software package. The vacuolar pH was estimated from a calibration curve that plotted the vacuolar pH of cells grown in APG media titrated to different pH values against the fluorescence intensities measured by the LSM700. Results and statistics were plotted using Graph Pad Prism 6.

### Cell Lines

The pQCXIP and IFITM3 plasmids and A549 cell lines were characterized previously. [21, 22] Briefly, A549 cells were grown in complete DMEM (Invitrogen #11965) with 10% FBS (Invitrogen). A549 cells were made by gamma-retroviral transducion with either an empty vector control or a vector containing the full-length human *IFITM3* cDNA. The cells were then selected with 2μg/mL puromycin in complete DMEM. Expression of IFITM3 was confirmed by Western blotting using an SDS-PAGE gel and an anti-IFITM3 antibody against the n-terminus of IFITM3 (Abgent #AP1153a).

### Lysotracker Red Staining

Lysotracker Red staining of A549 cells was done as described previously.[22] Briefly, A549 cells transduced with the empty vector or overexpressing IFITM3 were plated on coverslips and cultured for 4 hours in complete DMEM with either 20μM DMSO or SFN at 37°C. [16] For the last hour, Lysotracker Red DND-99 (Invitrogen) was added in the corresponding media to the cells. Cells were fixed with 4% PFA and stained with DAPI (blue). The coverslips were then imaged by a Leica SP-5 confocal microscope.

### SFN Survival Assay for Mammalian Cell Lines

Cells were plated in a 96-well plate at 8000 cells per well. They were then cultured with either 20μM DMSO or SFN in complete DMEM for 24 hours. [16] Cells were then fixed and stained with Hoechst and imaged by an IXM microscope. Meta-express software was used to count the number of cells indicated by DAPI staining.

## Results and Discussion

### Sulforaphane Inhibits the Growth of Wild Type Yeast Cells

Isothiocyanates have been used as antimicrobials, mainly for food preservation and plant pathogen control. [23] However, since sulforaphane (SFN), to the best of our knowledge at the time, had never been tested on yeast cells, we began by investigating whether the drug was able to inhibit the growth of wild type *Saccharomyces cerevisiae cells*. We plated ten-fold serial dilutions of wildtype cells from the PSY316 strain background on synthetic defined (SD) media with increasing concentrations of SFN (0-200 μg/ml). After two days of growth at 30°C, it was clear that SFN inhibited the growth of the strain (Figure 1A).

**FIGURE 1:**
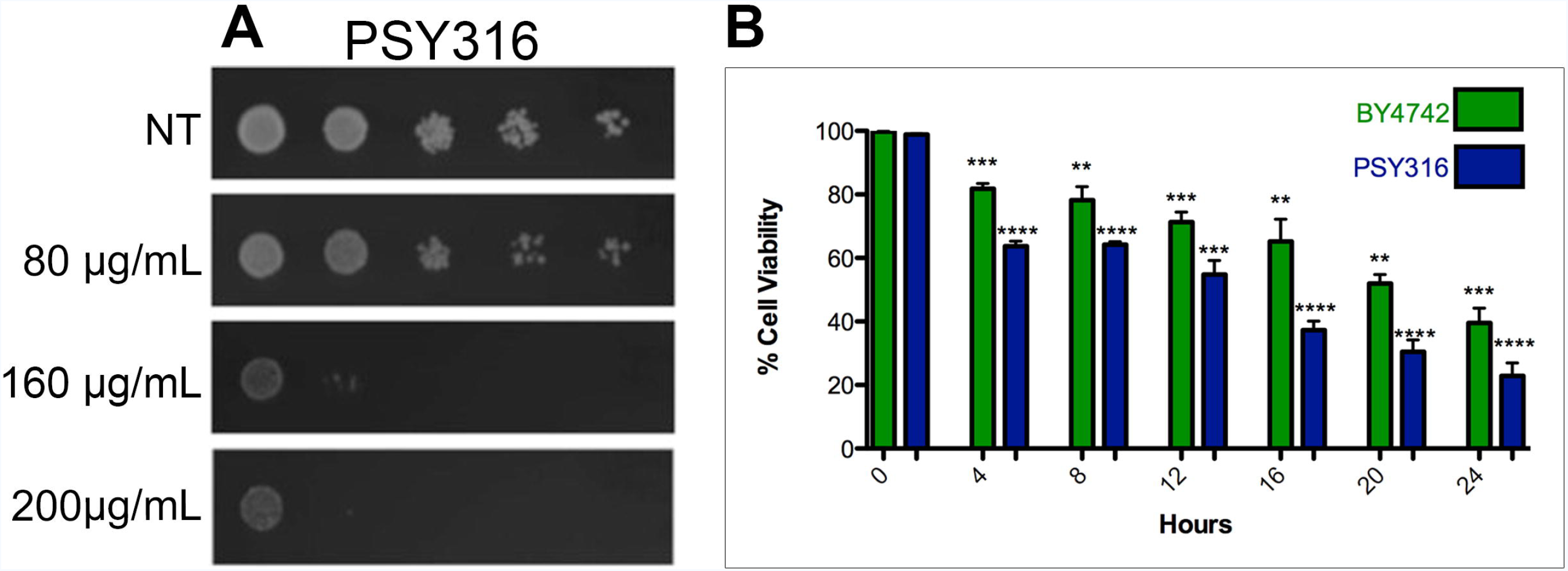
Sulforaphane Inhibits the Growth of Wild Type Yeast Cells. (A) Ten-fold serial dilutions of wildtype yeast cells from the PSY316 strain background were plated on synthetic defined media with increasing concentrations of SFN and allowed to grow at 30°C for two days. (B) Wildtype cells from both the BY4742 and PSY316 strain backgrounds were grown in synthetic defined liquid cultures containing 100μg/mL of SFN. The viability of the cells at the indicated time points was determined using propidium iodide as a vital stain. The difference in viabilities was deemed statistically significant by the Student’s t-test (p<0.05). Error bars indicate standard deviations for trials with at least three independent cultures.

Similar results were obtained when we measured the viability of the cells grown in liquid cultures containing 100 μg/ml SFN using propidium iodide as a vital stain, suggesting that the drug actually kills rather than merely inhibits yeast cell growth (Figure 1B). A parallel experiment with wildtype cells from the BY4742 strain background showed that the ability of SFN to trigger yeast cell death is not specific to a particular strain background (Figure 1B).

Given our laboratory’s wider interest in apoptotic-like cell death in yeast [24], we checked to see if SFN induces the characteristic hallmarks of this kind of cell death. We discovered that SFN-induced cell death neither generated reactive oxygen species (ROS), as determined by dihydroxyrhodamine 123 staining, nor was inhibited by the absence of oxygen (data not shown). Both are hallmark characteristics of apoptosis in yeast [25] suggesting that SFN-induced cell death is nonapoptotic in nature.

### A Genome-wide Screen Links Vacuolar Acidification to SFN’s Mechanism of Action

In order to better understand the mechanisms of action behind SFN-induced cell death, we undertook an unbiased genome wide screen to identify mutations that alter the cell’s sensitivity to SFN using the Saccharomyces cerevisiae knockout (YKO) library, a collection of 4,775 individual yeast strains in the BY4742 background, each of which contains a deletion of a single non-essential yeast ORF. Our initial experiments had revealed that 200μg/ml SFN significantly inhibits the growth of wildtype BY4742 yeast cells grown in 96-well liquid SD cultures for 48 hours, so we screened the YKO library for mutant BY4742 strains that were unable to grow under these conditions.

Each mutant strain was isolated by visually comparing 96-well plates with SFN to control plates without SFN, to identify wells that had little or no turbidity after 48 hours (Figure 2A). After screening the entire YKO library twice, we identified 311 mutant strains that consistently were unable to grow in liquid SD cultures containing 200μg/ml SFN after two days (Supplementary Table S1). Functional annotation utilizing gene ontology (GO) terms revealed that our screen had preferentially isolated mutants in genes involved in cellular metabolism, in the cell’s response to stress, and in the regulation of cell metabolism (Figure 2B). However, a cursory search through the Saccharomyces Genome Database (SGD) revealed that many, if not most, of these loss-of-function mutants are also sensitive to a wide range of other cellular insults and stresses suggesting that they may not be SFN-specific.

**FIGURE 2:**
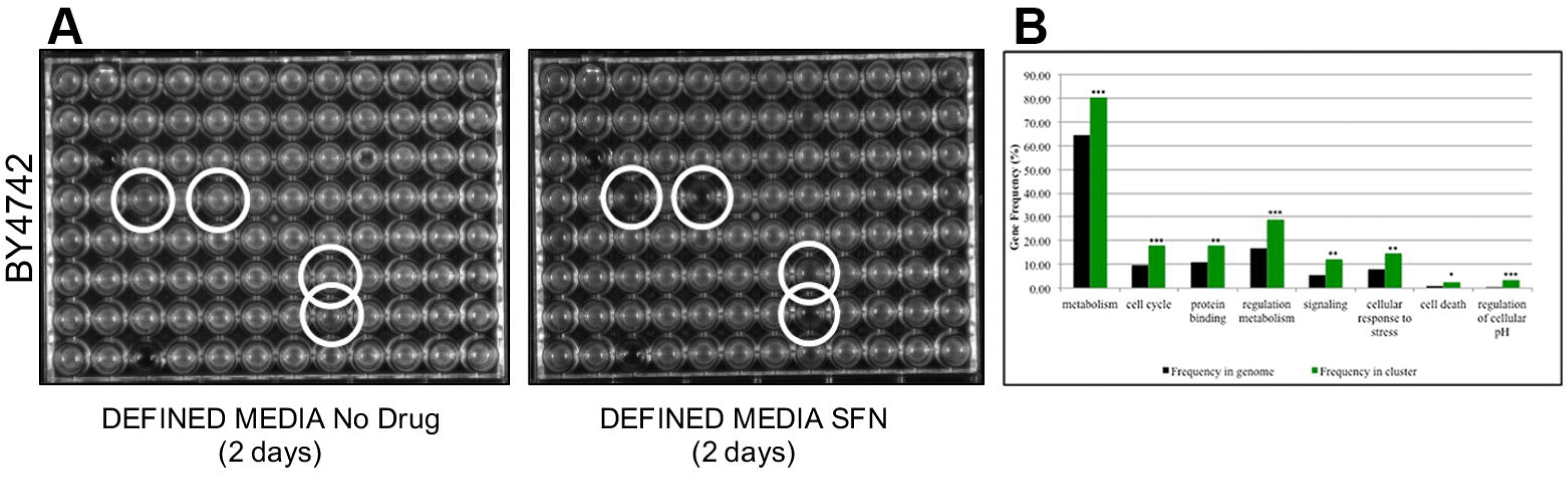
A Genome-wide Screen Links Vacuolar Acidification to SFN’s Mechanism of Action. (A) Seed cultures of individual yeast strains from the BY4742 knockout library were grown overnight at 30°C in 96-well plates in complete synthetic defined (SD) media and then transferred to media with and without 200μg/ml of SFN. Relative growth for SFN^S^ mutants was determined by visual inspection of the wells, comparing wells with drug with wells without drug, after they had been cultured at 30^°^C for 2 days. A representative pair of 96-well plates is shown. (B) Functional annotation utilizing gene ontology (GO) terms revealed that our screen had preferentially isolated mutants in genes involved in vacuolar function, especially in vacuolar acidification and/or pH regulation. Asterisks indicate statistical significance of the enrichment of ORFs identified in the screen as compared to their representation in the genome.

Intriguingly, however, we noticed that our SFN^S^ mutants were significantly enriched for genes involved in vacuolar function, especially in vacuolar acidification and/or pH regulation (Figure 2B). The vacuole has been implicated in the mechanism of action of numerous other drugs in yeast. [26-28] Our SFN^S^ vacuolar function deletion mutants included knockouts of *VMA1*, *VMA2*, and *VMA4*, which encode three of the subunits of the vacuolar H(+)-ATPase (V-ATPase) that is required for vacuolar acidification [29, 30]; knockouts of genes encoding the vacuolar fusion proteins, Vps41p, Vam3p, Vam6p, and Vam7p [31, 32]; and knockouts of the ergosterol biosynthesis proteins, Erg2p, Erg6p, and Erg24p. Notably, a previous study had linked genes involved in V-ATPase function, vacuolar fusion, and ergosterol biosynthesis to the vacuolar pH-stat of *Saccharomyces cerevisiae* [33], suggesting to us that the vacuole and especially the acidification of the vacuole may be linked to SFN function in yeast. A preliminary microarray and gene ontology (GO) analysis comparing ORFs upregulated in yeast cells grown in SFN as compared to cells grown in media containing other drugs such as benomyl, fluconazole, and paraquat also revealed an enrichment of vacuolar pH genes (data not shown).

### Lowering the Vacuolar pH Makes Yeast Cells Resistant to SFN-induced Cell Death

Because of the enrichment in our SFN^S^ screen of mutants linked to vacuolar acidification, we determined whether sulforaphane altered the vacuolar pH of the cell. Staining cells grown in SFN with the vacuole specific, pH-sensitive dye, 2,7’-bis (2-carboxyethyl)-5,6-carboxyfluorescein-acetoxymethylester (BCECF-AM), revealed that SFN significantly increases the vacuolar pH of two wildtype strains of different genetic backgrounds, making them more alkaline (Figure 3A and 3B).

**FIGURE 3:**
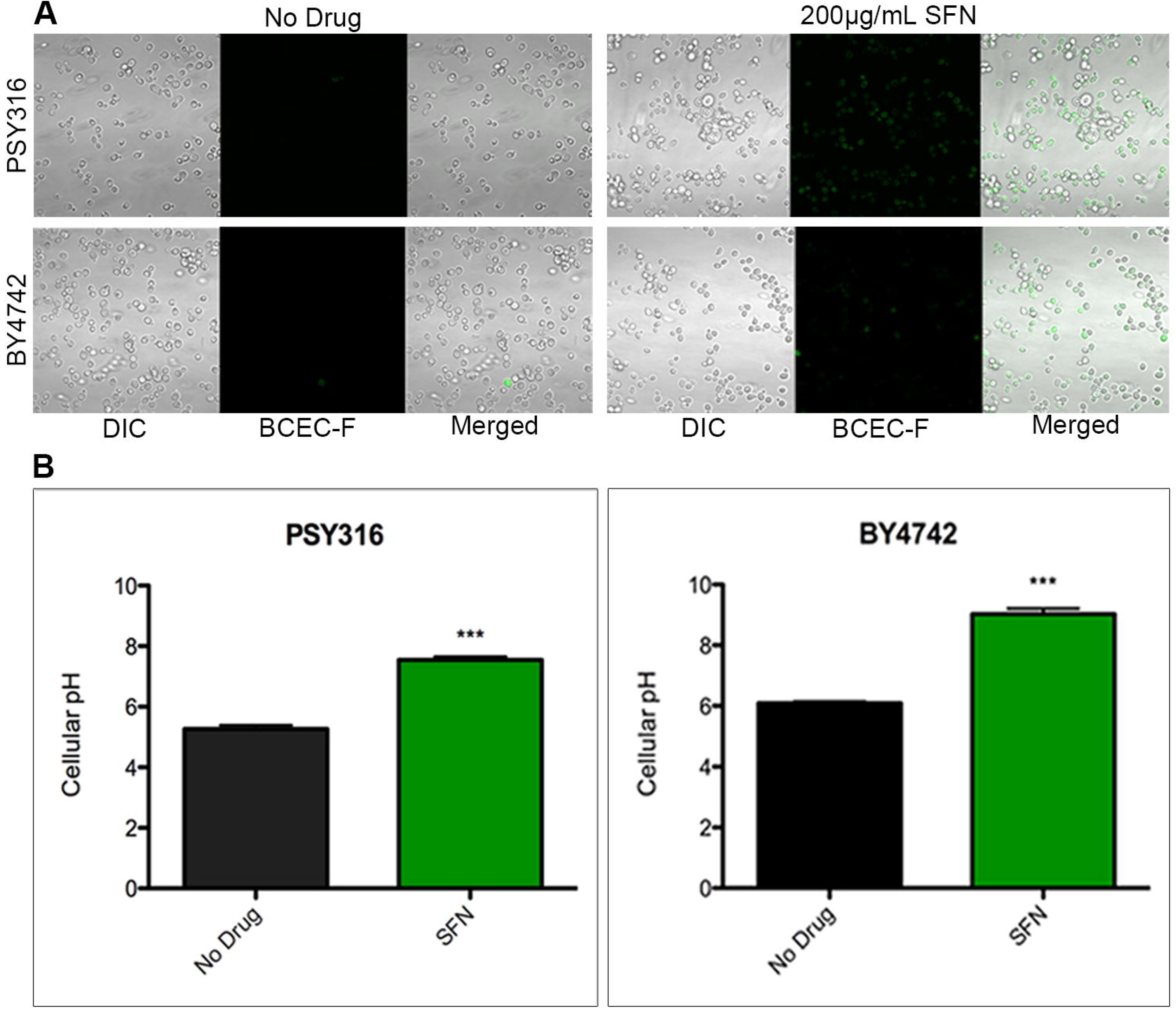
Sulforaphane Increases the pH of the Yeast Vacuole. (A) Wildtype cells from the BY4742 strain background were grown in synthetic defined liquid cultures containing 200μg/mL of SFN and were stained with the vacuole specific, pH-sensitive dye, BCECF-AM. Cells grown in SFN were significantly more fluorescent than their counterparts grown in media without drug. (B) The vacuolar pH of the cells imaged in Figure 3A was estimated from a calibration curve that plotted the vacuolar pH of cells grown in APG media titrated to different pH values against the fluorescence intensities measured by the LSM700. The difference in viabilities was deemed statistically significant by the Student’s t-test (p<0.05). Error bars indicate standard deviations for trials with at least three independent cultures.

From this observation, we hypothesized that SFN may trigger cell death by increasing the vacuolar pH of the yeast cell. To interrogate this possible mechanism of action, we sought to manipulate the vacuolar pH of the yeast cell to determine if this would alter the cell’s sensitivity to SFN. If SFN kills by increasing the pH of the yeast vacuole, we predicted that cells with more alkaline vacuoles than wildtype cells would be more sensitive to SFN because lower concentrations of the drug would more readily push cells beyond the threshold of alkalinity that triggers death. In contrast, we anticipated that cells with more acidic vacuoles would be more resistant to SFN than wildtype because it would take higher concentrations of the drug to push cells beyond a similar threshold.

The regulation of vacuolar pH in yeast is complex. [20] However, we took advantage of a battery of yeast vacuole acidification mutants, first identified by Brett et al. in a screen for genes involved in the vacuolar pH-stat in yeast, to see if we could discern a relationship between the pH of the yeast vacuole and the cell’s ability to grow on SFN plates. In this earlier screen, of the 107 mutants that displayed an aberrant vacuolar pH under more than one external pH condition, functional categories of transporters, membrane biogenesis, and trafficking machinery were significantly enriched.

Of the forty-six hyper-alkaline deletion strains determined by Brett et al. [20] to have more alkaline vacuoles than wildtype, eighteen (39%) were identified in our screen as SFN^S^ mutants. A Fisher exact test revealed that there was a statistically significant association between the two phenotypes of hyper-alkaline vacuoles and SFN^S^ (p<0.0001). On the other hand, of the seventy-seven hyper-acidic deletion strains known to have more acidic vacuoles than their wildtype counterparts, eleven (14%) were resistant to SFN (Figure 4; Table 1). These included deletions in genes involved in transcriptional and translational regulation (*RPL21B, RPS23B, RTF1, HAT1*) and sterol/lipid biogenesis (*SUR1*). A third of the SFN^R^ vacuolar hyper-acidic mutants were in genes of unknown function.

**FIGURE 4:**
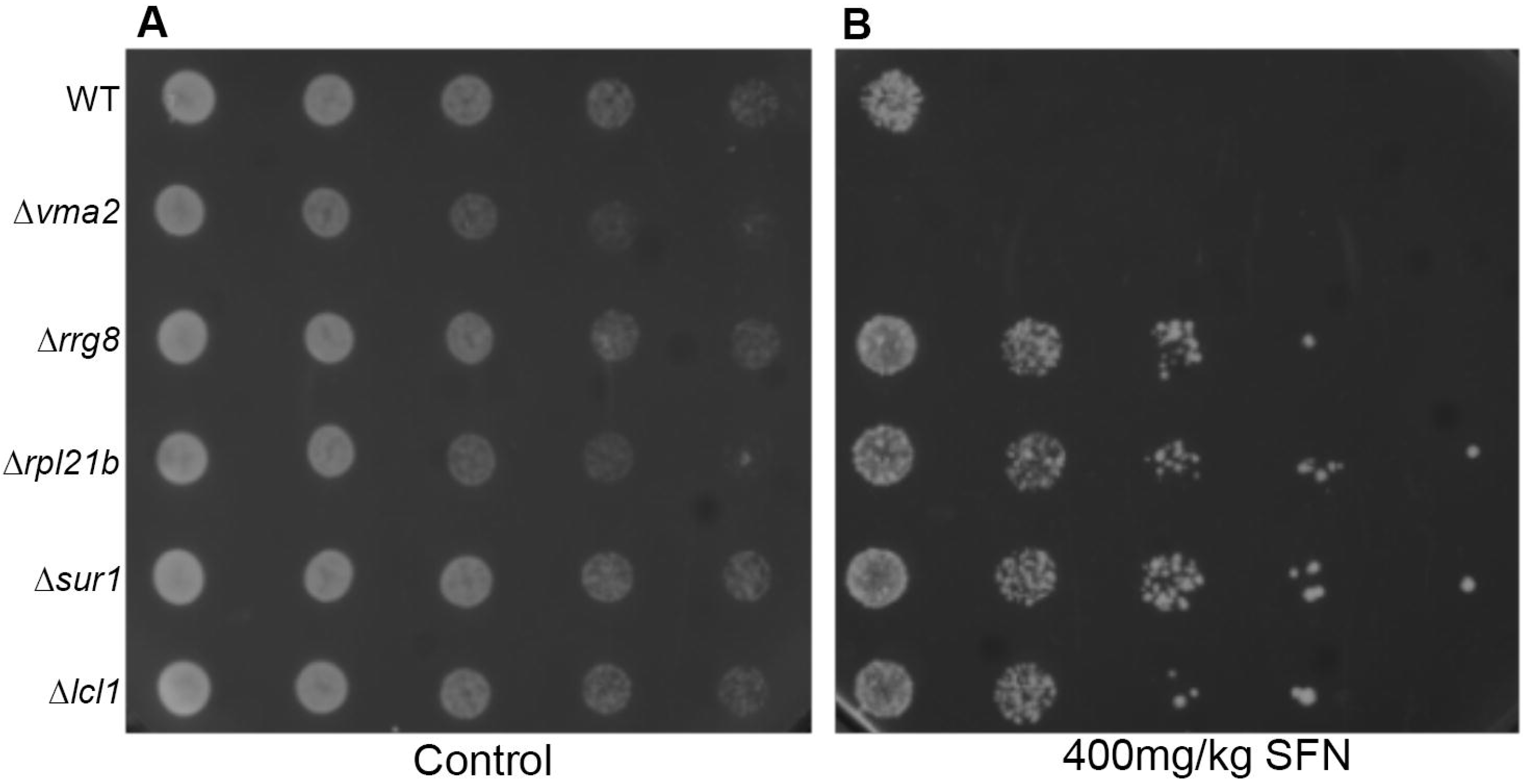
Mutations That Alter the Vacuolar pH of Yeast Cells Alter their Sensitivity to SFN. Ten-fold serial dilutions of yeast cells from the BY4742 strain background with mutations in genes known to regulate the pH of the yeast vacuole were plated on synthetic defined media with 400μg/mL SFN and allowed to grow at 30°C for two days. Deletions in genes known to increase vacuolar pH (*VMA2*) increased the sensitivity of cells to SFN while deletions in genes known to decrease vacuolar pH (*RRG8, RPL218, SUR1, and LCL1*) increased the resistance of cells to the drug.

**TABLE 1:** Eleven Genes Whose Deletions Are Known to Decrease Vacuolar pH Also Increase the Resistance of Yeast Cells to Sulforaphane.

| Gene Names (SFN<sup>R</sup>) |  |
| --- | --- |
| *COS12* | *RRG8* |
| *ECM23* | *RTF1* |
| *HAT1* | *SUR1* |
| *LCL1* | *TRM44* |
| *RPL21B* | *ULA1* |
| *RPS23B* |  |

It is not clear why only a subset of the vacuolar hyper-acidic mutants were SFN^R^, and we could not identify a common molecular explanation that would link them all to reveal SFN’s precise mechanism of action. However, given the complexity of the vacuolar pH-stat in yeast and the involvement of many of the hyper-acidic vacuolar genes in other physiological and metabolic pathways in the yeast cell, this should not be surprising. Nonetheless, this finding supports our claim that the mechanism of action of SFN-induced cell death involves the drug’s ability to increase the cell’s vacuolar pH. We still do not understand how SFN makes yeast vacuoles more alkaline and how an increase in vacuolar pH could trigger cell death. Though there is data that suggests that the V-ATPase promotes vacuolar membrane permeabilization (VMP) and nonapoptotic death in stressed yeast [34], we have discovered that SFN does not trigger this death-inducing mechanism: yeast cells grown in SFN and stained with FM4-64 to visualize their vacuolar membranes do not appear to undergo increased vacuolar fragmentation (data not shown).

### Sulforaphane’s Ability to Increase Vacuolar pH in Yeast is Drug Specific

As we have noted, the vacuole has been linked to the mechanisms of actions of a diversity of drugs and small molecules in yeast. [26-28] This raises the real possibility that an increase in the vacuolar pH is a generic response to drug insult in yeast. Recent studies suggest that the isothiocyanates phenethyl isothiocyanate (PEITC) and benzyl isothiocyanate (BITC), like SFN, can inhibit metastatic cell activity and migration. [35, 36] Therefore, to determine if SFN’s ability to increase the vacuolar pH is drug-specific, we checked to see if PEITC and BITC could similarly trigger an increase in vacuolar pH. If so, it would suggest that isothiocyanates in general and not SFN specifically kill cells via this mechanism.

As with SFN, we began by determining if PEITC and BITC could kill yeast cells in liquid culture. We found that the levels of cell death induced by 0.094μg/ml PEITC and 0.746μg/ml BITC were comparable to that triggered by 400μg/ml SFN (Figure 5A). However, in contrast with cells grown in SFN, yeast cells grown in PEITC and BITC did not increase their vacuolar pH as determined by BCEC-F staining (Figure 5B). This suggests that SFN’s mechanism of action in triggering cell death in yeast is distinct from the mechanisms used by two related isothiocynates, PEITC and BITC, to kill this simple eukaryote.

**FIGURE 5:**
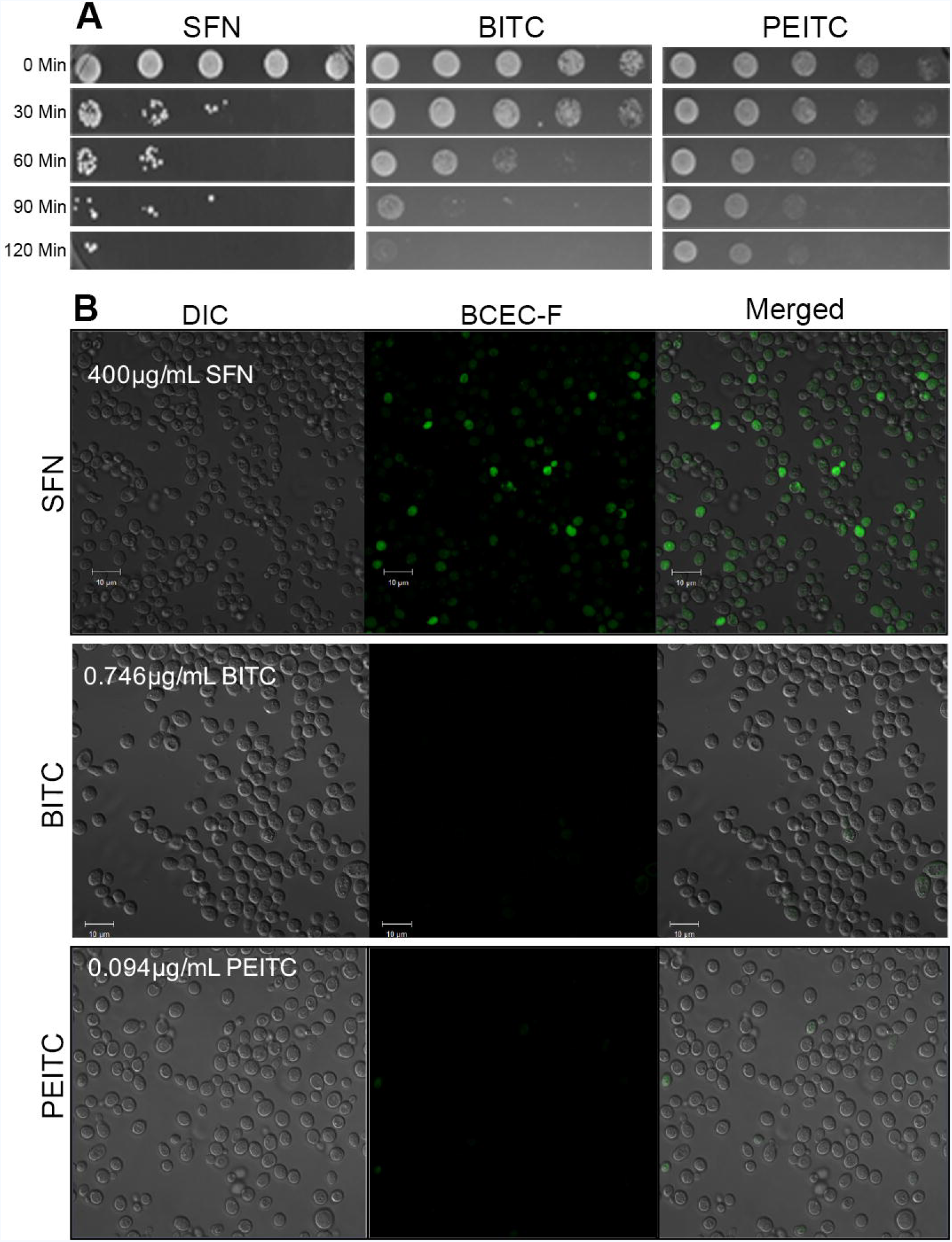
Sulforaphane’s Ability to Increase Vacuolar pH in Yeast is Drug Specific. (A) Ten-fold serial dilutions of wildtype yeast cells from the BY4742 strain background cultured in synthetic defined liquid cultures containing the indicated drugs, were plated on SD media and allowed to grow at 30°C for two days. (B) Wildtype cells from the BY4742 strain background were grown in synthetic defined liquid cultures containing the indicated drugs and were stained with the vacuole specific, pH-sensitive dye, BCECF-AM. Cells grown in SFN were fluorescent while their counterparts grown in media with the other drugs were not.

### Sulforaphane Increases the pH of Endosomes of Human A549 Cells

Given SFN’s well-studied ability to alter the physiology of mammalian cells, we visually examined A549 cells, a human alveolar adenocarcinoma cell line, cultured with SFN to determine if SFN’s mechanism of action in yeast cells is generally applicable. We discovered that A549 cells grown in media containing 40 μM SFN and the pH-sensitive dye, Lysotracker Red, show a decreased fluorescence as compared to cells grown in the absence of drug suggesting that they have more alkaline endosomes (Figure 6A). This suggests that SFN is able to increase the pH of both yeast vacuoles and mammalian lysosomes.

**FIGURE 6:**
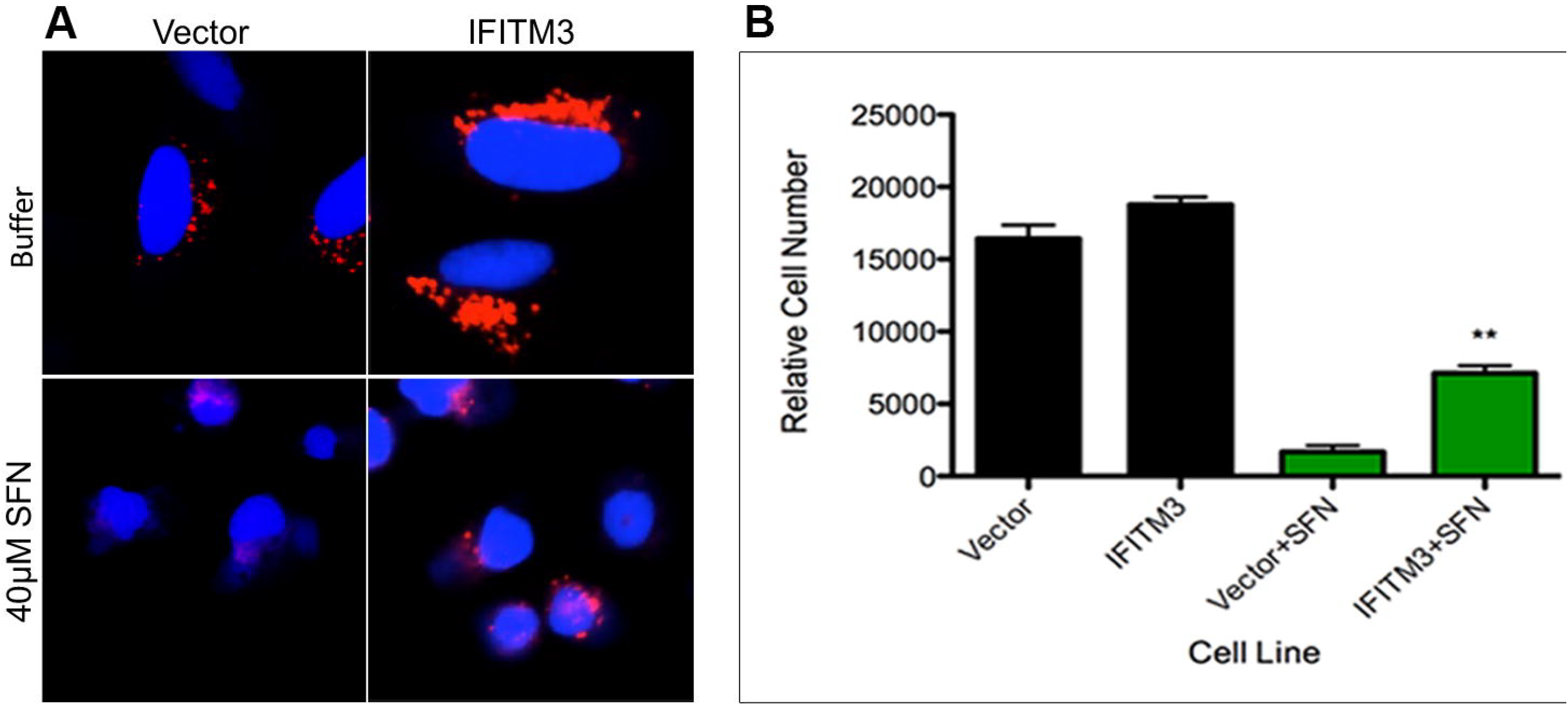
Sulforaphane Increases the pH of Endosomes of Human A549 Cells. (A) Cells from the A549 human alveolar adenocarcinoma cell line were cultured in media containing 40μM SFN and the pH-sensitive dye, Lysotracker Red. We also overexpressed the Interferon-inducible Transmembrane Protein 3 (IFITM3) protein that is known to enlarge the late endosomes and lysosomal compartments as well as increase their acidity in A549 cells. Cells grown with SFN are less positive for the dye than cells grown in the absence of the drug suggesting that they have more alkaline endosomes. (B) The viability of A549 cells with or without IFITM3 that had were cultured in media with or without SFN was determined by Hoechst staining. The difference in viabilities was deemed statistically significant by the Student’s t-test (p<0.05). Error bars indicate standard deviations for trials with at least three independent cultures.

Finally, in light of our findings that hyper-acidic yeast mutants are also resistant to SFN, we sought to make the lysosomes of mammalian cells more acidic to see if this too would in turn make them resistant to SFN. To do this, we overexpressed the Interferon-inducible Transmembrane Protein 3 (IFITM3) protein that is known to enlarge the late endosomes and lysosomal compartments as well as increase their acidity in A549 cells (Figure 6A) [22], and cultured the cells in SFN. We discovered that A549 cells overexpressing IFITM3 are relatively more resistant to SFN suggesting that lowering endosomal pH levels is also protective in higher eukaryotes (Figure 6B). It is a novel mechanism of action that should help us advance our understanding of sulforaphane’s chemopreventive and chemotherapeutic functions. Interestingly, it is a mechanism not unlike the mechanism of chloroquine which is known to increase the pH of the food vacuole of the plasmodium parasite, preventing hemoglobin degradation into a nontoxic digestible form. [37]

